# PIGSTI: a modular, reproducible pipeline for detecting species identity, pathogens, and microbes from animal palaeogenomic data

**DOI:** 10.64898/2026.09.01.748539

**Authors:** Louis L’Hôte, Catherine Butt, Áine Halpin, Luisa Sacristán, Valeria Mattiangeli, Pernille Bangsgaard, Lisa Yeomans, Melinda Zeder, Marjan Mashkour, Hossein Davoudi, Svend Hansen, Delphine Decruyenaere, Melissa Kennedy, Adeline Vautrin, Alisher Begmatov, Andrej B. Belinskiy, Amridin Berdimuradov, Gennadiy Bogomolov, Jacopo Bruno, Alexey Kalmykov, Jane McMahon, Jamal Mirzaakhmedov, Susan Pollock, Rocco Rante, Sabine Reinhold, Tobias Richter, Alisher Sandiboev, Eberhard Sauer, Laura Strolin, Hirofumi Teramura, Hugh Thomas, Jolijn A. M. Erven, Shigeki Nakagome, Daniel G. Bradley, Kevin G. Daly

## Abstract

Ancient genomics has enabled discovery of diverse pathogens across various time periods, host species, and material types. However, existing palaeogenomic pipelines predominantly focus on screening data from human hosts, or do not incorporate microbial screening methodologies. We present PIGSTI (Pathogen anImal Genome Sequence ToolkIt), a bioinformatic pipeline specifically designed for both the initial screening and subsequent detection of pathogens in shotgun sequencing data from ancient animal remains. PIGSTI’s integrated Snakemake workflow performs both host detection, genome mapping and pathogen identification, generating outputs suitable for population genetics and phylogenetic analyses. Testing on 952 newly sequenced and publicly available animal palaeogenomic datasets, we identified ∼15 ancient zoonotic and animal pathogens with high confidence, including the first documented case of *Rickettsia felis* and *Leptospira borgpetersenii* in an ancient animal. Our results demonstrate PIGSTI’s utility for screening pathogen diversity in ancient animal hosts and reconstructing historical host-pathogen relationships.

## Introduction

Ancient pathogen genomics has expanded rapidly in the past fifteen years, from the first draft genome of *Yersinia pestis* recovered from medieval remains (*1*) to the recent recovery of ancient RNA viruses (*2–4*). Research in this field has focused predominantly on human remains, revealing the evolutionary history of numerous human pathogens, including *Y. pestis* (*1*, *5–16*), HSV-1 (*17*), *Mycobacterium* species (*18*, *19*) (*19*) (*20*, *21*), (*22*, *23*), parvovirus19 (*24*), hepatitis B virus (*25–28*), measles virus (*2*), *Treponema pallidum* (*29–34*), variola virus (*35–37*), *Tarrenela forsythia* (*38*, *39*), *Plasmodium* species (*40–42*), *Salmonella enterica* (*43–45*), influenza viruses (*46*, *47*), and Rhinovirus (*4*). Despite this emphasis on human disease, several zoonotic pathogens and animal-specific pathogens have also been recovered from archaeological animal remains. These include *Brucella melitensis* from a sheep (*48*), sheeppox virus from archaeological sheep remains and codicological material (*49*, *50*), Marek’s disease virus from chicken bones (*51*), *Taenia hydatigena* from a sheep-associated calcified nodule (*50*), sheep-derived *Y. pestis* (*52*), and Megrivirus epengu and Rotavirus deltagastroenteritidis from penguin mummies (*53*). Secondary pathogens have also been reported in sheep, cattle and aurochs (*54*, *55*).

Ancient pathogen DNA recovery depends not only on the sampling material itself but also on the nature of the archaeological record more broadly. Animal remains are occasionally interred alongside humans as part of ritual practice, but far more often they represent butchery waste (*56*). Because such assemblages rarely comprise complete skeletons, opportunities to identify skeletal manifestations of chronic infection are correspondingly limited, with the exception of rare mass mortality assemblages, which can preserve substantial portions of an animal’s skeleton (51, 58). Faunal remains also differ from human remains in another aspect: as most derive from food waste, the animals in question were more likely to have been healthy, rather than diseased, at the time of death (*56*). Compounding these challenges, remains from multiple species are frequently commingled, and some skeletal elements, such as post-cranial bones from *Capra* and *Ovis*, cannot be reliably distinguished by morphological inspection alone (59, 60). DNA analysis of archaeozoological material therefore typically requires screening against multiple candidate host genomes to determine the most likely species of origin for each specimen. Competitive mapping approaches such as FastQ Screen (*60*) evaluate how reads are distributed across a set of reference genomes, while metagenomic classifiers such as Kraken2 (*61*) assign reads taxonomically without a predefined candidate list. Closely related taxa require dedicated methods: Zonkey, for example, compares low-coverage data against a panel equine genomes, discriminating equid species and identifying F1 hybrids from their intermediate ancestry (*62*).

In studies examining the genetic remains of ancient pathogens, host read identification and removal precedes any pathogen detection, which typically relies on metagenomic classification of the remaining non-host reads against modern reference databases. During this classification step, two complementary approaches are commonly followed. The first employs k-mer-based taxonomic classification, in which short subsequences (k-mers) from each read are matched against a reference database and assigned to the lowest common ancestor (LCA) of matching taxa. This strategy is implemented by the Kraken family of tools, including Kraken2 and KrakenUniq (*61*, *63*). KrakenUniq further improves confidence by reporting the number of unique k-mers assigned to each taxon. The second approach uses MALT (*64*), which directly aligns sequencing reads against a reference database. These alignments can then be evaluated using HOPS (*65*), which applies a series of authenticity criteria to assess the presence of ancient pathogens.

Several comprehensive workflows integrate these methods for ancient pathogen screening. EAGER (*66*, *67*) is a reproducible, portable and efficient pipeline that performs host read removal, metagenomic pathogen screening with HOPS, and downstream variant calling. The aMeta tool (*68*) combines KrakenUniq and HOPS, using KrakenUniq to identify candidate pathogens before restricting the subsequent HOPS analysis to those taxa. PALEOMIX (*69*) profiles microbial composition by mapping reads to a database of phylogenetically informative marker genes (*70*), an approach that becomes less reliable with highly fragmented ancient DNA. DNAharvester (*71*) screens host-unmapped reads against a user-supplied microbial reference database and produces authentication metrics, but is not primarily designed for microbial detection. Although such pipelines are increasingly commonly used, none combines accurate pathogen multi-methods identification with host identification, which is what animal palaeogenomic datasets require. EAGER includes host read removal but assumes that the host reference genome is already known, whereas aMeta does not include host read management. Because most ancient DNA studies focus on human remains, sequencing data are typically aligned directly to the human reference genome before pathogen screening. In contrast, analyses of archaeological animal remains require an initial host identification step before host read removal and pathogen screening can be performed, adding complexity to existing workflows and creating a barrier for those working with animal-derived palaeogenetic data.

Here, we present PIGSTI (Pathogen anImal Genome Sequence ToolkIt), a bioinformatic pipeline designed specifically for ancient animal genomics. PIGSTI identifies the host species from sequencing data using Fastq Screen (*60*), performs host genome alignment, and carries out metagenomic pathogen screening followed by high-resolution pathogen authentication. By default, PIGSTI uses KrakenUniq for pathogen screening together with the E-value filtering strategy described by Geuilil et al. (*72*), with optional HOPS-based validation. Candidate pathogens are subsequently aligned to reference genomes and evaluated using multiple authentication criteria to assess their authenticity. To demonstrate the performance of PIGSTI, we applied the pipeline to 952 ancient and modern genomic datasets from domestic livestock and wild animals, both published and newly generated, spanning approximately 75,500 years before present to modern specimens. Published datasets were drawn primarily from metAaRCive (version 26.05.08), a community-curated resource of metadata for published ancient animal genomes covering a wide range of taxa (*73*), from which we selected only livestock species and their wild relatives (*Bos*, *Ovis*, *Capra* and *Sus*), together with two further published datasets in which ancient microbes had previously been reported (*50*, *74*). We additionally explore the phylogenetic history of three disease-causing microbes recovered from this dataset: *Rickettsia felis*, *Leptospira borgpetersenii* and *Erysipelothrix rhusiopathiae*.

## Methods

### PIGSTI pipeline description

PIGSTI is a modular Snakemake workflow that integrates library quality control, host detection and mapping, and metagenomic and pathogen screening within a single run, making it well suited to the analysis of ancient DNA samples from a range of hosts (Figure 1). Reads first undergo adapter trimming and quality filtering. The pipeline then diverges into two parallel routes: a host-mapping route and a metagenomic screening route (Figure 1).

**Figure 1.**
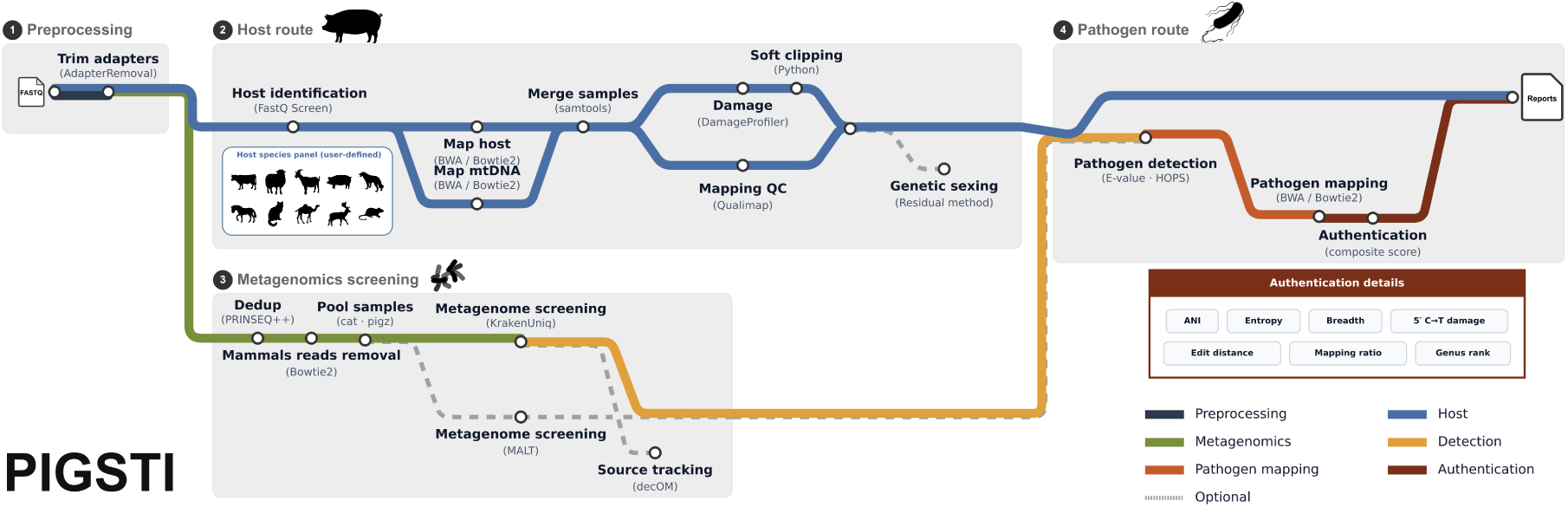
Schematic overview of the PIGSTI analysis workflow. The flowchart shows the main subworkflows of PIGSTI: preprocessing, host identification and mapping, metagenomic screening, and pathogen detection and authentication. Each subworkflow is separated by grey outlines, with arrows indicating the direction of data flow between them. Line colours correspond to each subworkflow, as indicated in the legend, and dashed lines indicate optional steps.

Host route: host identification is performed using FastQ Screen (*60*), which conducts competitive mapping of adapter-trimmed reads against a user-defined panel of host genomes. PIGSTI assigns the host species based on two criteria: the species receiving the highest total number of mapped reads, and the species with the highest number of reads mapping uniquely to a single genome (“one hit, one genome”). This screening step is initially run on a subsample of the data; if fewer than 50 reads meet the one-hit-one-genome criterion, as can occur in samples with very low endogenous DNA content, the analysis is repeated on the full dataset. PIGSTI additionally flags human contamination, which can arise from extensive handling, such as parchment (*49*).

In the host route, quality-trimmed reads are aligned to the mitochondrial and nuclear genomes of the identified host using a short-read aligner with parameters suited to damaged ancient DNA (*75*, *76*). Low-quality alignments are removed and PCR duplicates are marked and discarded. From the resulting BAM files, PIGSTI calculates endogenous DNA content (defined as [number of final mapped reads / number of raw reads] × 100), genome coverage, and, depending on the host species and reference genome used, can determine the genetic sex of the individual based on the ratio of reads aligning to each chromosome versus chromosome length (*77*).

Metagenomic route: in parallel, the metagenomic route characterizes the broader microbial community and screens for pathogens or specific microbes of interest. Because metagenomic abundance estimates are sensitive to duplicate reads, and because metagenomic analyses are computationally demanding, exact-duplicate reads are first removed from the adapter and quality-trimmed reads. The de-duplicated reads are then mapped against a user-defined panel of potential host and contaminant genomes (including human) to remove host-derived and contaminant sequences prior to downstream analysis. Using the resulting unmapped reads, PIGSTI can optionally perform metagenomic source-tracking with decOM (*78*), an approach particularly relevant to applications such as dental calculus studies (*79*).

Following source-tracking, PIGSTI performs pathogen screening using KrakenUniq (*63*) and, optionally, MALT via the HOPS pipeline (*65*). KrakenUniq is well suited to identifying low-abundance taxa, as it reports the number of unique k-mers assigned to reads alongside an estimate of breadth of coverage. For each species detected, PIGSTI calculates an E-value, following Guellil and colleagues (*72*), defined as E = (K/R) × C, where K is the unique k-mer count, R is the read count, and C is the estimated breadth of coverage. Using a user-defined pathogen list, PIGSTI flags as a potential candidate any species passing E-value thresholds, which can be set either as defaults or at the level of individual pathogens. This allows, for example, more stringent filtering for opportunistic commensals such as *Escherichia coli*, while applying more permissive thresholds for pathogens such as poxviruses. Samples in which a pathogen is flagged via E-value and/or HOPS (if enabled) are then mapped to the corresponding pathogen genome using the same parameters as in the host-mapping step.

For each mapped pathogen, PIGSTI performs a series of post-mapping analyses and generates a score out of 10 (or 13 if HOPS is enabled), based on the following metrics, described in detail below: KrakenUniq read count, E-value, entropy, edit-distance decay (damaged and non-damaged read subsets), postmortem damage, average nucleotide identity, breadth ratio, mapping ratio, and genus ranking.

We now present a definition of criteria used in pathogen detection and post-mapping authentication. Some of these criteria are drawn directly from Pathopipe, developed by Sikora and colleagues to screen large-scale human datasets (*80*).

KrakenUniq read count: the clade-level read count from the KrakenUniq report (i.e., reads assigned to a taxon plus all of its descendant taxa, as distinct from taxReads, which reflects only reads assigned to the terminal node). A minimum clade read count of 50 is used as the default threshold.

E-value: following Guellil et al (*72*), the E-value is calculated for each candidate taxon as E = (K/R) × C, where K is the number of unique k-mers assigned to the taxon, R is the number of reads assigned to the taxon, and C is the estimated k-mer coverage (genome breadth) reported by KrakenUniq. Elevated E-values reflect reads distributed across a larger genomic fraction with proportionally unique k-mer support. The default detection threshold is E > 0.001.

Average nucleotide identity (ANI): ANI is defined as the mean percentage of identical nucleotides across all local alignments between two genomic sequences. Here it is approximated directly from read-mapping data as ANI ≈ (1 − mismatches/bases mapped) × 100, using mismatch and mapped-base counts derived from samtools stats. Reduced ANI may reflect damage-related substitutions, mapping to a distantly related reference, or misclassification of reads originating from a related species. The default threshold is >96.5%.

Relative entropy: a measure of how evenly mapped reads are distributed across the reference genome, adapted from Sikora and colleagues. Read start positions are binned into non-overlapping windows of 100 bp and 1,000 bp, and the Shannon entropy of the resulting distribution is calculated and normalized to the maximum entropy attainable under a uniform distribution: H_rel = H / H_max = [−Σ p_i log₂(p_i)] / log₂(N_windows), where p_i denotes the proportion of read starts in window i. Values approaching 1 indicate coverage consistent with a genuine, evenly distributed pathogen signal, whereas low values are consistent with localized clustering indicative of contamination or mismapping. Default thresholds, following Sikora and colleagues (*80*), are ≥0.9 for bacteria/archaea and ≥0.7 for viruses.

Breadth ratio: a measure of coverage evenness, adapted from Sikora and colleagues., calculated as the ratio of observed to expected genome breadth: B_observed / B_expected, where B_expected = 1 − e^(−mean depth) is the breadth predicted under a Poisson model of uniform read distribution at a given sequencing depth. Values approaching 1 indicate coverage consistent with even mapping across the genome; values well below 1 indicate patchy or clustered mapping, potentially reflecting cross-mapping from a related taxon. The default threshold is ≥0.8.

Edit-distance decay: edit distance describes the number of mismatches between a read and the reference it is mapped to. Reads are split into damaged and non-damaged subsets based on the presence of terminal deamination (5′ C→T and/or 3′ G→A within the terminal five aligned bases), following the same logic used by HOPS (*65*). For each subset, a composite Decay Quality Score (range 0–1) is calculated incorporating: monotonicity of read count decline across mismatch bins (weight 0.35), the ratio of reads at edit distance 0 relative to edit distance 1 (0.25), the proportion of reads at edit distance 0 (0.15), a penalty applied where the modal edit distance is not 0 (0.10), and the rate (0.10) and R² (0.05) of an exponential decay fit to the mismatch distribution. Default pass thresholds are 0.65 (damaged reads) and 0.55 (non-damaged reads).

Postmortem damage: the frequency of C→T substitutions at the terminal 5′ position of aligned reads, estimated using DamageProfiler (*81*). A minimum threshold of 0.01 is applied.

Mapping ratio.: a measure of concordance between KrakenUniq classification count and reference alignment, calculated as the number of mapped reads (BWA/Bowtie2) divided by the number of KrakenUniq-assigned reads. The default threshold is ≥0.5.

Genus rank: the rank of a candidate taxon relative to all other species of the same genus reported by KrakenUniq, ordered by descending clade read count. A rank of 1, i.e., the dominant genus-level hit, is required to award a detection point, reducing the likelihood of ambiguous assignment among closely related taxa.

HOPS criteria: when HOPS is enabled, PIGSTI additionally aligns reads for each candidate pathogen using MALT and evaluates three further authentication criteria via MaltExtract, contributing up to 3 additional points to the score (for a maximum of 13).

These criteria assess, for each positive hit, the decline of the edit-distance distribution, the presence of characteristic terminal aDNA damage patterns, and the proportion of damaged reads among those with edit distance zero, following the authentication framework of HOPS (*65*).

#### PIGSTI pipeline - tool and parameter details

We provide an overview of each of the tools used across the PIGSTI pipeline.

1. Pre-processing

● Adapter trimming: AdapterRemoval v2 (*82*); Cutadapt (*83*) for single-end libraries.
● Host species identification: FastQ Screen (*60*) with Bowtie2 (*75*).
2. Host / mtDNA mapping and post-processing

● Alignment: BWA aln (*76*) or Bowtie2 (*75*).
● Mapping quality filtering, duplicate marking: Samtools markdup (*84*) or Picard’s (http://broadinstitute.github.io/picard) MarkDuplicate function.
● Damage profiling: DamageProfiler (*81*).
● Mapping QC: Qualimap2 (*85*).
● Terminal 4 bases soft-clipping: PIGSTI custom python script softclip_mod.py.
● Genetic sexing (optional): implemented for cattle, goat, sheep, and dog.
3. Metagenomic screening

● Dereplication: PRINSEQ++ (*86*).
● Mammalian host-read removal: Bowtie2 (*75*) against a multi-host index, retaining unmapped reads.
● Metagenomic classification: KrakenUniq (*63*).
● Optional metagenomics classification: HOPS/MALT/MaltExtract (*65*).
● Optional source tracking: decOM (*78*).
4. Pathogen detection, mapping, and authentication

● Candidate selection: KrakenUniq read counts and Guellil E-value against a user-defined pathogen spreadsheet (in addition to HOPS, if enabled).
● Pathogen alignment: BWA aln (*76*) or Bowtie2 (*75*).
● Mapping quality filtering, duplicate marking: Samtools markdup (*84*) or Picard’s (http://broadinstitute.github.io/picard) MarkDuplicate function.
● Damage profiling: DamageProfiler (*81*).
● Mapping QC: Qualimap2 (*85*).
● Authentication: KrakenUniq read count, E-value, entropy, edit-distance decay, damage, ANI, breadth ratio, mapping ratio, and genus rank (plus three additional MaltExtract-based criteria if HOPS is enabled).

#### Simulation of ancient pathogen reads

To assess PIGSTI’s detection performance across a range of pathogens and pathogens abundances, we generated paired-end mock ancient metagenomes in which a single pathogen genome was spiked into a fixed host and environmental background. Each library was built to a total of 12 million reads, of which 10% comprised endogenous sheep DNA from Menteşe6, a west Turkish Neolithic sample previously found positive for *B. melitensis* (*48*, *87*). The remaining reads represented microbial background drawn from Menteşe5, a sample from the same site previously found negative for pathogen DNA. Within each mock metagenomic community, 60 simulated samples were generated by spiking in simulated reads from one of ten pathogens (*B. melitensis*, *Coxiella burnetii*, *E. rhusiopathiae*, *L. borgpetersenii*, Sheeppox virus, *T. pallidum*, Variola virus, *Y. pestis*, *Mycobacterium leprae*, and *Bacillus anthracis*) at one of six abundance levels (80, 345, 1,500, 6,400, 28,000, and 120,000 reads). The proportion of environmental background reads was adjusted at each abundance level to keep total library size constant.

Pathogen-derived paired-end reads were simulated using NGSNGS (*88*), with read lengths drawn from a normal distribution (mean 60 bp, s.d. 20 bp). Post-mortem damage was modelled using the NGSNGS Briggs damage model (-m Illumina,0.024,0.36,0.68,0.0097; (*89*)), and stochastic divergence from the modern reference genome was introduced at a mutation rate of 0.0001 substitutions per site per generation.

### Analysis of ancient livestock dataset

Genetic data of 153 archaeozoological specimens from 32 archaeological sites are newly reported here. Details of each archaeological site are provided in the “Description of archaeological sites for newly reported material” section following the Discussion.

#### Laboratory methods

All laboratory work was carried out in dedicated ancient DNA facilities at Trinity College Dublin. Work surfaces were routinely cleaned with dilute (0.5%) sodium hypochlorite, researchers wore full-body personal protective equipment at all times and changed gloves frequently. Consumables were UVC-irradiated immediately prior to use, and all milling and drilling equipment was decontaminated between samples with bleach, isopropanol and UVC exposure.

#### Sampling

Teeth were sectioned along the cementoenamel junction and the root halved longitudinally. Material was collected from the pulp chamber and the interior of the root using a dental drill, yielding 12–104 mg of powder per sample.

For petrous bone and other skeletal elements, the surface was first cleaned with a dental drill and a Dremel saw was used to remove a subsample from the densest region of the element. Subsamples were pulverised in decontaminated stainless-steel mill jars with grinding balls using a MixerMill at 30 Hz, with repeated cycles until fully powdered, yielding 54–130 mg per sample.

#### DNA extraction

DNA was extracted following the protocol of Mattiangeli et al. 2023 (*90*), with the pre-treatment adjusted to the substrate.

Teeth were not bleach-washed. Powder was subjected to a 15-minute pre-digestion in 1 ml 0.5 M EDTA at 37 °C, after which the supernatant was discarded. Petrous portions and other skeletal elements received a dilute sodium hypochlorite wash (0.5% for 15 minutes, followed by three 1 ml washes in H₂O) before a 30-minute EDTA pre-digestion under the same conditions. Foetal bones and teeth were not bleach-treated.

The remaining pellet was digested overnight at 37 °C with frequent agitation in 1 ml of extraction buffer (per sample: 17 μl N-lauroylsarcosine, 20 μl 1 M Tris-HCl, 940 μl 0.5 M EDTA and 13 μl 50 U/ml proteinase K; the mastermix was UVC-irradiated for 30 minutes before proteinase K was added). Following centrifugation at 17,000 g for 10 minutes, the supernatant was purified on Roche High Pure Extender Assembly columns with 13 ml of modified PB buffer (per sample: 0.42 ml 3 M sodium acetate, 0.33 ml 5 M sodium chloride and 12.25 ml PB buffer, Qiagen). DNA was eluted in 50 μl EBT.

#### Library preparation and sequencing

Double-stranded libraries were prepared following Meyer and Kircher 2010 (*91*). For each sample, one library was built without uracil-DNA glycosylase treatment, using 12–16 μl of input DNA, in order to preserve authentic damage patterns; all remaining libraries were treated with 5 μl UDG (New England Biolabs, M5505L) at 37 °C for 1 hour prior to construction. Libraries were dual-indexed, amplified for 12 PCR cycles, and shotgun-sequenced on an Illumina NovaSeq X platform with 150 bp paired-end reads. Sequencing statistics and metadata for each indexed amplification are given in Table S2 in the extended data.

#### Applying PIGSTI

Shotgun libraries from published and newly produced samples were adapter-and quality-trimmed using AdapterRemoval v2 for paired-end data or Cutadapt for single-end data, using the parameters described above. Host species was identified against reference genomes for cow, dog, human, horse, sheep, pig, goat, deer, cat, rat, and camel (ARS-UCD1.2, ROS_Cfam_1.0, GRCh38, EquCab3.0, ARS-UI_Ramb_v2.0, Sscrofa11.1, ASM170441v1, mCerEla1.1, Fca126_mat1.0, Rrattus_CSIRO_v1, CamDro3); for computational efficiency, FastQ Screen was run only on the initial read subset, without full-dataset rescreening of low-one-hit samples. Samples with less than 20 reads assigned to one read one genome were considered as uncertain hosts. For previously published datasets, host species were already known and host mapping was disabled; for newly generated datasets, host nuclear and mitochondrial genomes were mapped using Bowtie2 in end-to-end sensitive mode, and genetic sexing was not performed.

For metagenomic screening, reads were depleted against the a multi-mammalian host Bowtie2 index (cow, dog, human, horse, sheep, pig, goat, deer, cat, and camel), and unmapped, pooled reads were classified using KrakenUniq against the NT_2020_microbes database (NCBI non-redundant NT records from June 2020 that includes all microbial organisms (archaea, bacteria, viral, fungi, protozoa, parasitic_worms) plus human reference genome and a few complete eukaryotic genomes, as implemented in aMeta (*68*). Due to computational limitations, HOPS/MALT and decOM were not used in this run.

Candidate pathogens were selected using the E-value and read-count thresholds described above (read-count thresholds varied by pathogen; due to the size of this dataset, the full list is provided as Table S1 in the extended data). Candidates were mapped to their reference genomes with Bowtie2 in end-to-end sensitive mode and authenticated using the ten criteria described above.

#### Phylogenetic placement

To evaluate whether low-coverage ancient detections could be reliably placed within a modern reference phylogeny, we used PathPhynder (*92*), which places ancient samples onto trees built from modern and high-coverage ancient genomes using a reference tree and a corresponding VCF built against a single-chromosome reference genome. Because the standard implementation (v1.23) discards a sample from a branch once the number of alternative alleles inconsistent with that branch exceeds a fixed tolerance, an approach well suited to taxa with limited diversity but poorly suited to highly mutable, diverse organisms and to low-coverage ancient BAM files of variable depth, where SNP informativeness scales with coverage rather than remaining constant. We therefore modified PathPhynder v1.23 to use a proportional rather than fixed maximum tolerance for alternative alleles (modified source code provided in the extended data), applying a tolerance proportion of 0.45 throughout this study. For each taxon, a modern reference phylogeny was reconstructed with Snippy (https://github.com/tseemann/snippy), using the resulting core.aln to build a maximum likelihood tree in IQ-TREE2 (*93*) with ModelFinder (best-fitting substitution model, 1,000 bootstrap replicates), and the corresponding core.vcf as the SNP collection for PathPhynder placement.

For *Rickettsia*, this reference panel comprised the genome dataset published by Rifkin et al. (*94*); as the *R. felis* genome is distributed across multiple contigs, we built a concatenated reference by joining contigs with 100-N spacers, then ran Snippy against it using the 31 modern genomes together with the ancient samples STK240 and MC72713, whose cleaned, mammalian-depleted reads were remapped with BWA aln (parameters as above), yielding 346 reads for MC72713 and 1234 for STK240; the resulting midpoint-rooted tree was used for placement (nucleotide substitution model TVM+F+ASC+R2 as determined by ModelFinder and 1,000 ultrafast bootstrap replicates). Pathphynder detected 48659 representative SNPs.

For *L. borgpetersenii*, complete genomes were downloaded from RefSeq via NCBI Datasets (*95*), and, as for *Rickettsia*, a concatenated reference was built from the genome’s contigs. An initial phylogeny built with *Leptospira interrogans* as an outgroup left too few SNPs shared across the alignment for reliable placement of the ancient genome Kaz4 (see below), so the tree was rebuilt with IQ-TREE2 (nucleotide substitution model TVM+F+ASC as determined by ModelFinder and 1,000 ultrafast bootstrap replicates) excluding *L. interrogans* and midpoint-rooted instead. Kaz4’s cleaned, mammalian-depleted reads were then remapped with BWA aln (parameters as above) to this concatenated reference, which served as the Snippy reference (59102 representative SNPs detected by Pathphynder).

For *E. rhusiopathiae*, complete *E. rhusiopathiae* genomes together with an *Erysipelothrix tonsillarum* reference genome were downloaded from RefSeq using NCBI Datasets (*95*). Snippy was run against the modern *E. rhusiopathiae* reference genome using the modern genomes together with the high-coverage ancient genome Galicia3, and a maximum likelihood phylogeny was reconstructed in IQ-TREE2 (nucleotide substitution model TVM+F+ASC+R2 as determined by ModelFinder and 1,000 ultrafast bootstrap replicates) and with *E. tonsillarum* as an outgroup (17923 representative SNPs detected by Pathphynder).

#### Virulence gene coverage analysis

The presence and coverage of 56 characterized E. rhusiopathiae virulence genes were assessed using whole-genome sequencing reads mapped to the reference contig NZ_LR134439.1. Gene coordinates were extracted from the reference genome annotation, and per-gene coverage metrics were calculated for each sample using Samtools (*96*) and BEDtools (*97*). A locus was scored as present when per-gene read coverage reached ≥ 75% breadth for samples sequenced to > 3× depth (Galicia3) or ≥ 50% breadth for samples sequenced to < 3× depth (Dublin2, YG13, YG01).

## Results

### Testing PIGSTI on simulated data

Having developed the PIGSTI pipeline (described above), we sought to test it on simulated metagenomic data featuring pathogen-derived reads. To explore how this approach deals with cases of pathogen DNA being present at differing degrees, we simulated ancient metagenomic datasets containing reads from ten pathogens at varying abundance (*B. melitensis*, *C. burnetii*, *E. rhusiopathiae*, *L. borgpetersenii*, sheeppox virus, *T. pallidum*, variola virus, *Y. pestis*, *M. leprae* and *B. anthracis*). We simulated six pathogen-read abundance levels for all ten pathogens within metagenomic datasets of 12 million reads each, of which 10% were endogenous (sheep) host DNA and the remainder represented microbial background; pathogen-derived reads ranged from 80 to 120,000 per dataset. Out of the ten pathogens, two, *M. leprae* and *B. anthracis*, were not detectable with PIGSTI using KrakenUniq alone as the metagenomic screening step. Although *M. leprae* itself was not detected, PIGSTI identified *M. avium* at low confidence within the same simulated dataset (full simulation results provided as Table S3 in the extended data). Of the remaining eight detectable pathogens, five (*C. burnetii*, *E. rhusiopathiae*, *L. borgpetersenii*, sheeppox virus, variola virus) were detected at every abundance level tested, while the remaining three (*B. melitensis*, *T. pallidum*, *Y. pestis*) required at least 345 reads to be reliably detected.

All detectable taxa achieved the maximum authentication score (10/10) once pathogen abundance reached approximately 1,500 reads (Figure 2A). At lower abundances (80–345 reads), some pathogens received reduced scores (8–10/10), failing most often on relative entropy (lowest observed: 0.42) and breadth ratio (lowest observed: 0.50) (Figure 2B). To retain true-positive low-abundance samples while maintaining specificity, we relaxed the breadth ratio and relative entropy thresholds relative to the default scoring criteria, but observed that all eight detectable pathogens showed edit-distance decay in the absence of aDNA damage of at least 0.7, enabling us to use this as a stringent filter. The resulting high-confidence filtering thresholds applied to subsequent empirical analyses were: breadth ratio ≥ 0.5, relative entropy ≥ 0.45, genus ranking = 1, edit-distance decay in the absence of aDNA damage ≥ 0.7, ANI > 96.5%, and mapping ratio > 0.5 (Figure 2B).

**Figure 2.**
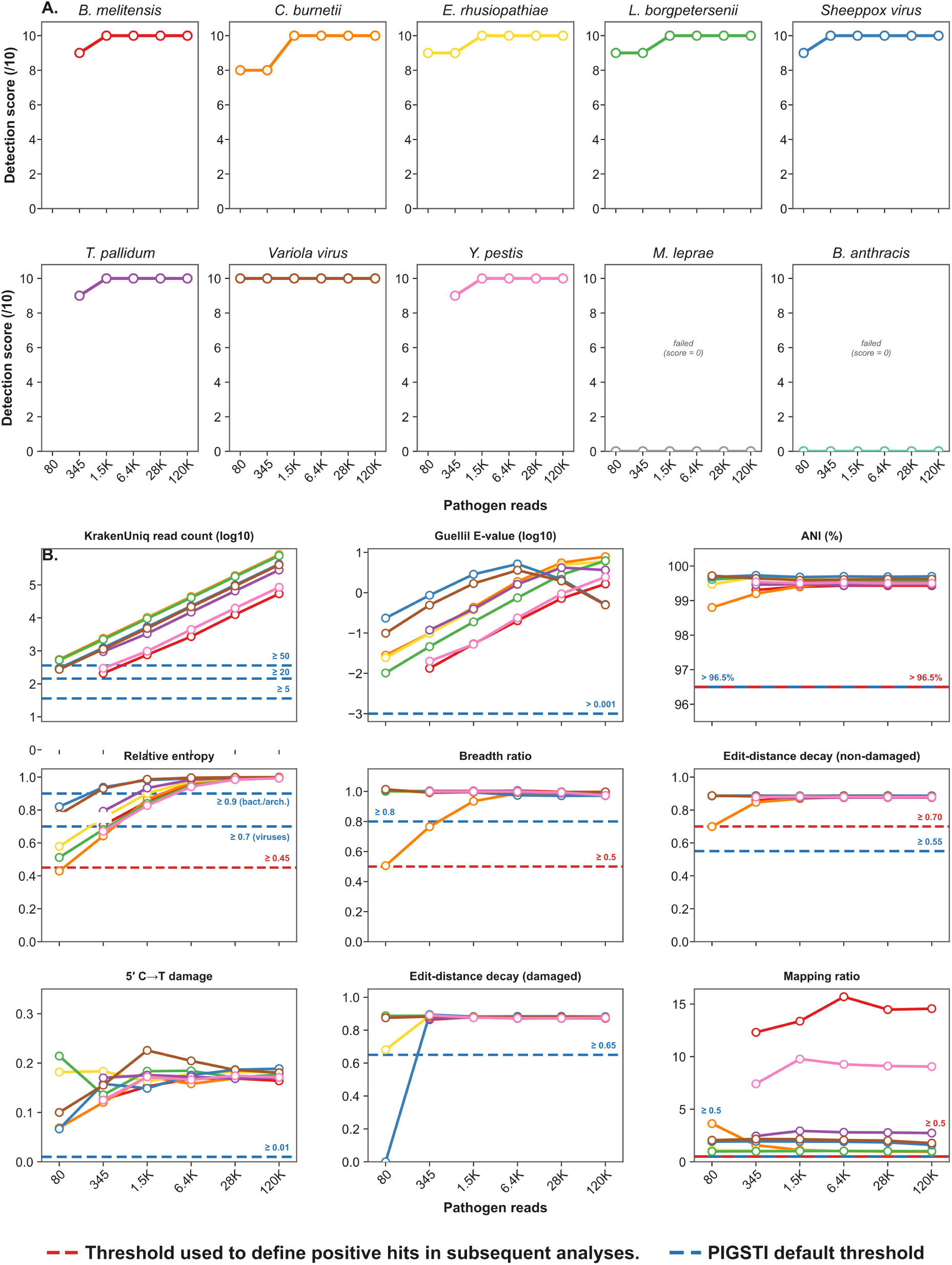
PIGSTI benchmarking on simulated metagenomic data. (A) Detection score (out of 10) for each of ten spike-in pathogens across six pathogen-read abundance tiers (80, 345, 1,500, 6,400, 28,000, and 120,000 reads). *M. leprae* and *B. anthracis* failed detection (score = 0) at all abundance tiers and are excluded from panel B. (B) PIGSTI screening metrics versus pathogen-read abundance for the eight successfully detected pathogens. Coloured lines correspond to individual pathogens presented in panel A. Blue dashed lines indicate PIGSTI default scoring thresholds initially applied to the simulated dataset. Red dashed lines indicate the modified thresholds subsequently used to define positive hits in empirical data, based on the simulation results (breadth ratio ≥ 0.5, relative entropy ≥ 0.45, edit-distance decay ≥ 0.7, ANI > 96.5%, mapping ratio > 0.5). Genus ranking is excluded from the plot as it was always = 1 across all detections.

### Applying PIGSTI to empirical data

To develop a high-level overview of disease-causing microbes in animals in the past, we used PIGSTI, with the empirically tested parameters described above, to screen 952 ancient and modern domestic and wild animal genomic datasets, spanning approximately 75,500 years before present to the modern day. Our data comprised 799 previously published datasets, taken directly from metAaRCive (version 26.05.08) (*73*) and restricted to livestock species and their wild relatives (*Bos*, *Ovis*, *Capra* and *Sus*), 2 published datasets in which ancient microbes had previously been reported (*50*, *74*) and 153 newly generated datasets. The published datasets originated from a range of sample materials, including soft tissues (n = 8), petrosal bones (n = 261), teeth (n = 42), long bones (n = 34), other skeletal elements (n = 21), parchments (n = 8), and samples for which material information was unavailable (n = 425). Host species represented in the published datasets included *Sus* (n = 354), *Ovis* (n = 165), *Capra* (n = 146) and *Bos* (n = 134), among others. The 153 newly generated datasets consisted of long bones (n = 71), teeth (n = 57), petrosal bones (n = 19) and other skeletal elements (n = 6). Species identification of the newly generated specimens assigned 34 samples to *Ovis*, 34 to *Capra*, 7 to *Bos*, 3 to *Sus* and 1 to *Camelus*, while 74 samples remained unresolved (see Methods for species identification criteria). Full metadata for all screened samples are provided as Table S3 in the extended data.

PIGSTI pathogen screening was run on a panel of zoonotic and animal-specific pathogens using KrakenUniq combined with E-value filtering. Of these, 1,023 microbe hits passed the E-value criteria across the dataset. Of the 1,023 pathogen hits, 109 passed these high-confidence filters (relative entropy ≥ 0.45, genus ranking = 1, edit-distance decay (no damage) ≥ 0.70, breadth ratio ≥ 0.50, ANI > 96.5% and mapping ratio > 0.50) across 89 samples from 952 screened (Figure 3A). These 89 samples span roughly 13,000 years to the present, and comprise 33 petrosal bones, 25 teeth, 10 long bones, 5 parchment folia, 1 other skeletal element, 2 soft tissues and 13 samples with missing material information. They represent 37 *Ovis*, 22 *Capra*, 20 *Bos*, 6 *Sus* and 4 samples of an uncertain host. Geographically, detections span a broad range from Iceland to the Eurasian Steppe (Figure 3A).

**Figure 3.**
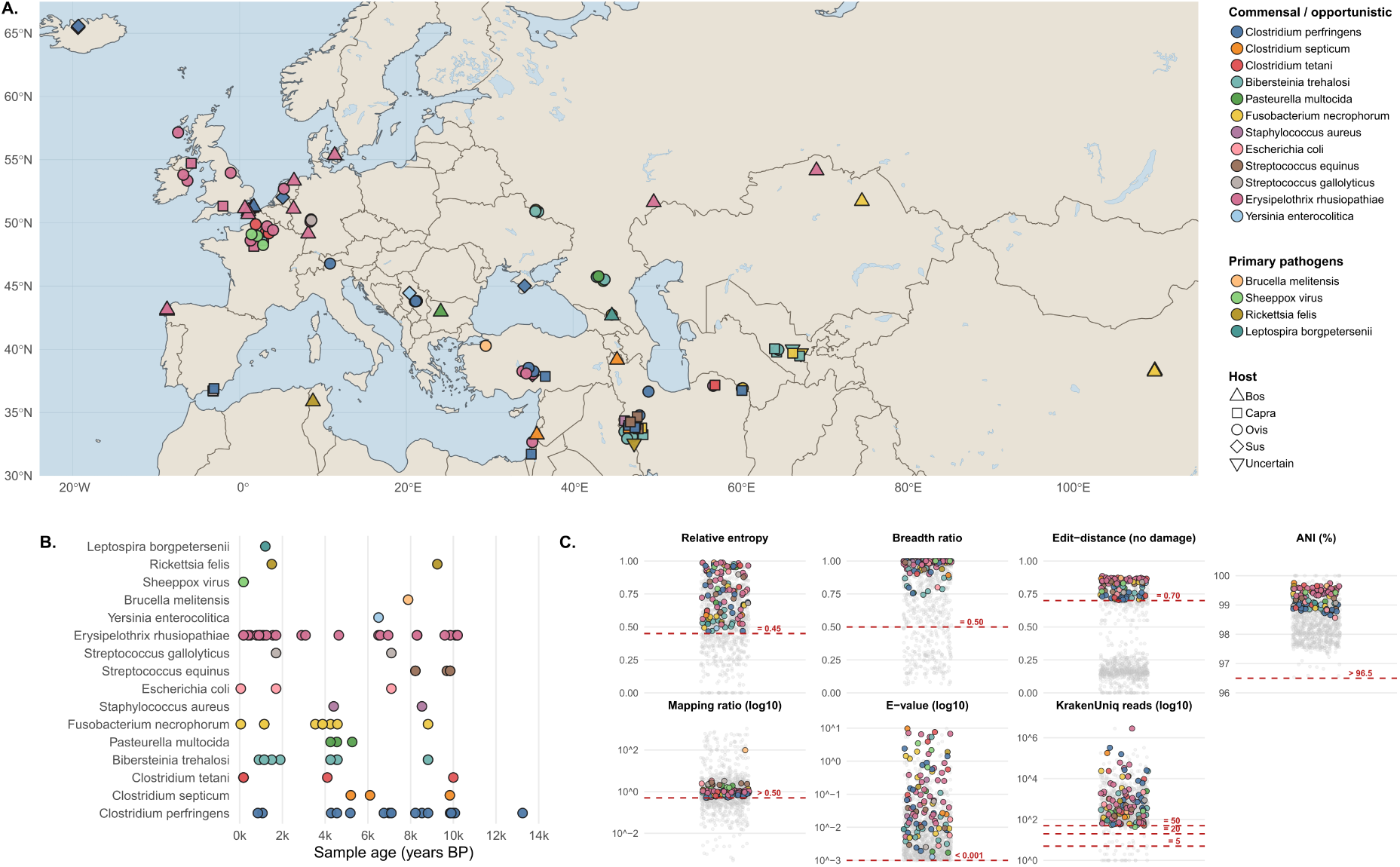
Spatial, temporal and authentication profiles of pathogen detections. (A) Map showing the geographic distribution of pathogen detections among the 952 ancient and modern animal datasets screened with PIGSTI. Points are coloured by pathogen and shaped by host species (n = 107 detections from 88 samples with coordinates; two authenticated detections from one sample without coordinates are omitted). (B) Temporal distribution of the 100 authenticated pathogen detections with a reported sample age, plotted by sample age (years Before Present) (109 authenticated detections in total). (C) Authentication metrics for pathogen detections (n = 1,023 pathogen detections). Coloured points denote the 109 detections that passed the full filter set; grey points denote detections that did not. Detections were retained if relative entropy ≥ 0.45, genus ranking = 1, edit-distance decay (no damage) ≥ 0.70, breadth ratio ≥ 0.50, ANI > 96.5% and mapping ratio > 0.50. Red dashed lines also show reference thresholds for E-value (< 0.001) and KrakenUniq read counts (5, 20 and 50).

We note a potential uneven distribution of authenticated detections across host species and sample materials relative to the screened cohort (Table 1). Host representation among the 89 positive specimens differed significantly from expectation (chi-square test, 5 degrees of freedom: χ²₅ = 47.1, P = 5.5 × 10⁻⁹): *Ovis* accounted for 42% of samples with authenticated detection (37/89) but only 21% of those screened (×2.0), with *Capra* (×1.3) and *Bos* (×1.5) also overrepresented. In contrast, *Sus* contributed 7% of positive specimens (6/89) despite comprising 38% of the screened cohort (×0.2). Sample materials were similarly non-random (χ²₄ = 66.9, P < 10⁻¹²): dental tissues accounted for 28% of positive specimens (25/89) compared with 10% of those screened (×2.7), while cranial and dental tissues together accounted for 65% of positive specimens (58/89). This uneven distribution holds when two studies reporting only sheepox virus-positive specimens (*49*, *50*, *74*) are excluded (χ²₅ = 41.0, P = 9.4 × 10⁻⁸; χ²₄ = 43.8, P = 7.0 × 10⁻⁹).

**Table 1.**
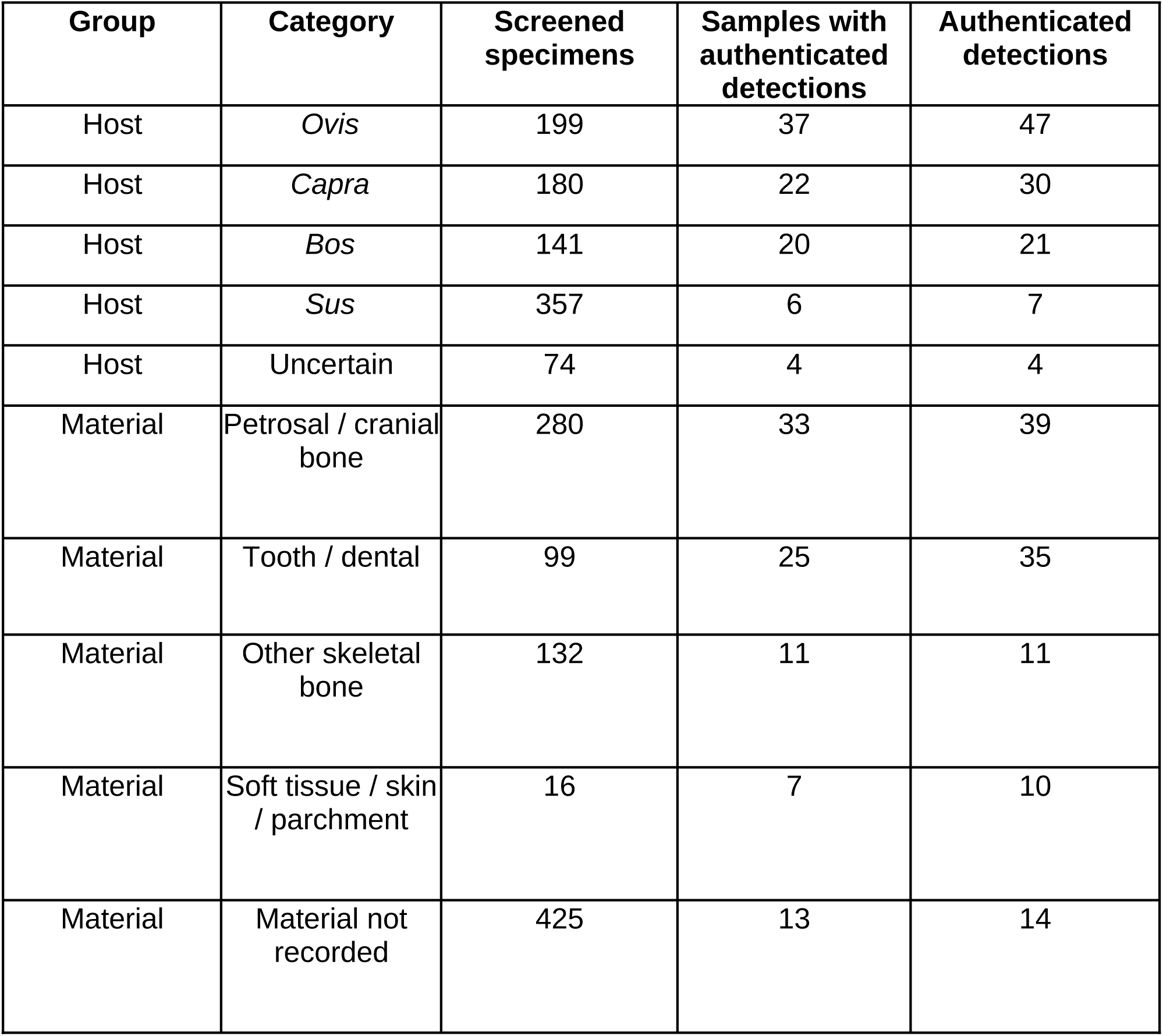
Host species and sample material among authenticated PIGSTI pathogen detections.

We grouped the detected pathogens into functional categories reflecting their typical ecology and transmission. The underlying screening results, with and without filtering, are provided as Tables S5, S6 and S7 in the extended data.

**Commensal/opportunistic pathogens.**

This category comprises organisms that can persist in healthy animals but become pathogenic under certain conditions, or that are otherwise present in the environment and become infectious following wounds, ingestion, or other routes of entry. It includes *Clostridiales* species such as *Clostridium perfringens* (N = 18, depth of coverage range: 0.0008–3.7051×), *Clostridium septicum* (N = 3, depth of coverage range: 0.0022–3.5904×), and the tetanus agent *Clostridium tetani* (N = 5, depth of coverage range: 0.0015–0.4408×). *Clostridium septicum* is implicated in braxy (or bradsot) (*98*), and both *C. septicum* and *C. perfringens* are implicated in enterotoxaemia, an economically damaging acute diseases of sheep and cattle (*98–100*), while *C. tetani* causes tetanus following wound contamination (*101*, *102*). We also detected Gram-negative, facultatively anaerobic coccobacilli of the family *Pasteurellaceae*, agents of pasteurellosis and bovine/ovine respiratory disease: *Bibersteinia trehalosi* (N = 15, depth of coverage range: 0.0010–0.5823×) and *Pasteurella multocida* (N = 3, depth of coverage range: 0.0045–0.1428×). These organisms remain leading causes of respiratory disease, such as pasteurellosis, and of sudden mortality in modern sheep and cattle husbandry, frequently acting opportunistically following stress, transport, or viral co-infection (*103*). We further identified *Fusobacterium necrophorum* (N = 11, depth of coverage range: 0.0011–0.5175×), commonly associated with footrot and other necrotic infections in livestock (*104*).

We additionally detected *Staphylococcus aureus*, known to cause mastitis in livestock (N = 2, depth of coverage range: 0.0156×–0.0681×) (*105*), and *E. coli*, certain strains of which are responsible for watery mouth disease in young lambs (*106*) and intestinal disease in adults (*107*) (N = 3, depth of coverage range: 0.0187×–0.1004×). Members of the *Streptococcus bovis*-type subgroup, classic inhabitants of the rumen capable of causing ruminal acidosis and systemic lactic acidaemia (*108*), were also detected, consistent with previous reports from ancient small ruminant specimens (*54*). These included *Streptococcus equinus* (N = 6, depth of coverage range: 0.0033–0.0351×) and *Streptococcus gallolyticus* (N = 2, depth of coverage range: 0.0043–0.0439×). We also recovered *Yersinia enterocolitica* from a pig sample (BLT104) (N = 1, depth of coverage: 0.0007×); *Y. enterocolitica* is usually a ubiquitous, commensal organism in pigs but can become pathogenic in humans (*109*). Finally, we detected *E. rhusiopathiae*, an opportunistic, Gram-positive, non-spore-forming, facultative intracellular pathogen with a broad host range, notably responsible for swine erysipelas (*110*). This pathogen has previously been reported in numerous ancient datasets, notably from humans (*54*, *55*, *111*); and we detect it here in three animals (Galicia3, Galicia1 and Win1) also reported by de-Dios et al. (*55*). Overall, *E. rhusiopathiae* was detected in 33 animal samples, with coverage ranging from 0.0017× to 67.7123×.

**Obligate pathogens.**

The second pathogen group comprises organisms that are exclusively infectious and not found in healthy individuals, representing a major threat to livestock health. We recovered a previously published *B. melitensis* genome (N = 1, depth of coverage: 0.3907×), the agent of the most prevalent bacterial zoonosis worldwide (*112*), and the major livestock pathogen sheeppox virus in 3 previously published samples (depth of coverage range: 0.0520×–0.2746×) (*49*, *50*). We identified two *R. felis* genomes: one in a Tunisian cattle sample (STK240) dated contextually to the 5th century CE (*113*), and one in newly generated data from MC72713, a metapodial bone from Ali Kosh, Iran (∼9,250 BP) lacking a resolved genetic host assignment. *Rickettsia felis* is an obligate intracellular bacterium of the transitional group of *Rickettsia* species and the causative agent of flea-borne spotted fever, with the cat flea *Ctenocephalides felis* as its principal vector (*114*). A genome of *R. felis* has previously been recovered from the remains of a child dated to approximately 2,000 years ago in southern Africa (*94*). Although no direct evidence of cattle infection has previously been reported, cattle have been shown to carry potentially infected fleas (*115*). Finally, we detected *L. borgpetersenii* in Kaz4, a medieval cattle sample from Georgia (*116*) (N = 1, depth of coverage: 0.0011×), an obligate pathogenic spirochaete and agent of leptospirosis in livestock, which can cause abortion, infertility, and decreased milk production in cattle (*117*). It is also an important global zoonosis, causing an estimated 50,000 human deaths annually (*118*). Evidence of *L. borgpetersenii* has also been detected in Neolithic Sweden (∼5000 BP) from human remains (*80*).

### Confirming pathogen identity through phylogenetic placement

To confirm that the putative pathogens identified by our screening pipeline corresponded to the taxa assigned by PIGSTI, we validated the placement of three candidates, *R. felis*, *L. borgpetersenii*, and *E. rhusiopathiae*, using maximum likelihood phylogenies built from modern and high-coverage ancient genomes (depth of coverage >10×), onto which we placed low-coverage ancient samples using PathPhynder. Table 2 displayed samples that we attempted to place into phylogenies:

**Table 2.** Metadata of samples considered likely pathogen detection.

| Sample | Pathogen | Host | Tissue | Approximate sample age (BP) | Country of origin | Site of origin |
| --- | --- | --- | --- | --- | --- | --- |
| STK240 | *Rickettsia felis* | *Bos* | Missing | 1500 | Tunisia | Althiburos |
| MC72713 | *Rickettsia felis* | *Uncertain* | Long bone | 9250 | Iran | Ali Kosh |
| Kaz4 | *Leptospira borgpetersenii* | *Bos* | Missing | 1200 | Georgia | Dariali/Tamara Fort |
| AA401 | *Erysipelothrix rhusiopathiae* | *Sus* | Missing | 1065 | Iceland | Skagaþjörður |
| AL725 | *Erysipelothrix rhusiopathiae* | *Sus* | Missing | 10200 | Turkey | Aşıklı Höyük |
| Asikli4 | *Erysipelothrix rhusiopathiae* | *Ovis* | Long bone | 10200 | Turkey | Aşıklı Höyük |
| Asikli7 | *Erysipelothrix rhusiopathiae* | *Ovis* | Petrous bone | 9600 | Turkey | Aşıklı Höyük |
| Asikli8 | *Erysipelothrix rhusiopathiae* | *Ovis* | Petrous bone | 9860 | Turkey | Aşıklı Höyük |
| Bed4 | *Erysipelothrix rhusiopathiae* | *Bos* | Petrous bone | 10108 | Germany | Bedburg-Königshoven |
| Carrick3 | *Erysipelothrix rhusiopathiae* | *Capra* | Long bone | 394 | Ireland | Carrickfergus |
| Dublin1 | *Erysipelothrix rhusiopathiae* | *Ovis* | Petrous bone | 1250 | Ireland | Moynagh Crannog |
| Dublin2 | *Erysipelothrix rhusiopathiae* | *Ovis* | Petrous bone | 900 | Ireland | Fishamble Street |
| Enkhuizen 1 | *Erysipelothrix rhusiopathiae* | *Ovis* | Petrous bone | 3080 | Netherlands | Enkhuizen |
| Falconeri2 | *Erysipelothrix rhusiopathiae* | *Capra* | Tooth | -38 | Missing | Born at the MHNH zoo, France |
| Frankfurt1 | *Erysipelothrix rhusiopathiae* | *Ovis* | Petrous bone | 1700 | Germany | Frankfurt-Heddernheim |
| Galicía1 | *Erysipelothrix rhusiopathiae* | *Bos* | Petrous bone | 8297 | Spain | O Courel, Galicia |
| Galicía3 | *Erysipelothrix rhusiopathiae* | *Bos* | Long bone | 8311 | Spain | O Courel, Galicia |
| Hungate1 | *Erysipelothrix rhusiopathiae* | *Ovis* | Petrous bone | 800 | United Kingdom | Hungate |
| Hxh2 | *Erysipelothrix rhusiopathiae* | *Bos* | Petrous bone | Missing | Germany | Herxheim |
| Kilpheder 2 | *Erysipelothrix rhusiopathiae* | *Ovis* | Petrous bone | 900 | United Kingdom | Kilpheder |
| MDVC564 | *Erysipelothrix rhusiopathiae* | *Ovis* | Petrous bone | 6950 | France | Menneville-Derrière-Le-Village, Picardie |
| Potterne1 | *Erysipelothrix rhusiopathiae* | *Capra* | Petrous bone | 2900 | United Kingdom | Potterne, Wiltshire |
| RF003 | *Erysipelothrix rhusiopathiae* | *Ovis* | Tooth | 177 | France | Louvres "Rue de Bouteiller" |
| RF009 | *Erysipelothrix rhusiopathiae* | *Ovis* | Tooth | 177 | France | Louvres "Rue de Bouteiller" |
| ROS002 | *Erysipelothrix rhusiopathiae* | *Bos* | Astragalus | 4637 | Kazakhstan | Roshchinskoe |
| VEM185 | *Erysipelothrix rhusiopathiae* | *Sus* | missing | 6500 | United Kingdom | Durrington Walls |
| Var1 | *Erysipelothrix rhusiopathiae* | *Bos* | Petrous bone | 6580 | Russia | Varfolomeevka |
| Walie1 | *Erysipelothrix rhusiopathiae* | *Capra* | Tooth | 0 | Missing | Born at the MNHN zoo, France |
| Win1 | *Erysipelothrix rhusiopathiae* | *Bos* | Missing | 1556 | Netherlands | Winsum-Bruggeburen |
| YG01 | *Erysipelothrix rhusiopathiae* | *Ovis* | Parchment | 500 | United Kingdom | Canterbury |
| YG13 | *Erysipelothrix rhusiopathiae* | *Bos* | Parchment | 500 | United Kingdom | Canterbury |
| YG14 | *Erysipelothrix rhusiopathiae* | *Bos* | Parchment | 945 | United Kingdom | Canterbury |
| YG15 | *Erysipelothrix rhusiopathiae* | *Bos* | Parchment | 945 | United Kingdom | Canterbury |
| YG16 | *Erysipelothrix rhusiopathiae* | *Bos* | Parchment | 945 | United Kingdom | Canterbury |
| Yoqneam 1 | *Erysipelothrix rhusiopathiae* | *Ovis* | Petrous bone | 1130 | Israel | Tel Yoqne'am |
| Zea1 | *Erysipelothrix rhusiopathiae* | *Bos* | Petrous bone | Missing | Denmark | Lundby I |

Given the limited number of complete modern *R. felis* assemblies, we used the same dataset as Rifkin et al. (*94*) and built a maximum likelihood phylogeny of 31 genomes spanning different *Rickettsia* species, including the previously reported ancient BBayA strain. The ancient samples STK240 and MC72713 reported here were placed with high confidence on the branch leading to *R. felis* (Figure 4A): at the *R. felis* node, 103 of 104 informative SNPs supported the placement of STK240 and 22 of 24 supported MC72713, confirming their taxonomic identity. The low number of reads recruited to the genomes (n = 346 reads for MC72713 and n =1234 for STK240) precluded further downstream analysis of this strain.

**Figure 4.**
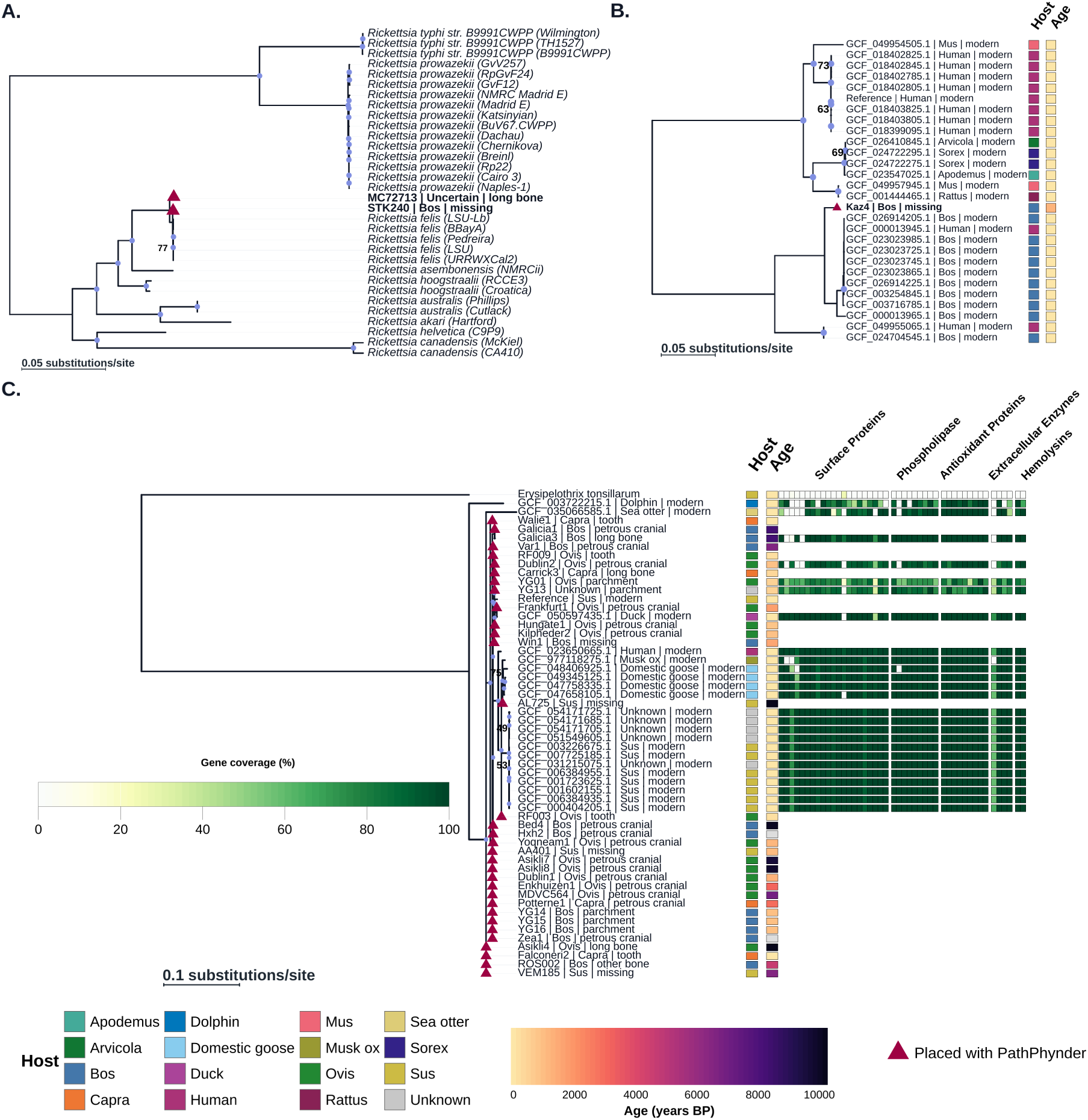
Phylogenetic placement of ancient pathogen detections. Maximum likelihood phylogenies of modern and high-coverage ancient genomes, with low-coverage ancient detections placed using PathPhynder (red triangles; placed sample labels in bold on panels A and B). Blue circles mark nodes with bootstrap support >80%; numeric labels show support ≤80%. Adjacent Host and Age annotation heatmap summarise host taxon and sample age; grey cells in the Age column indicate missing age information for ancient samples. (A) Midpoint-rooted phylogeny of 32 *Rickettsia* genomes showing PathPhynder placement of the ancient samples STK240 and MC72713 on the branch leading to *R. felis*, alongside the previously reported ancient BBayA strain. (B) Midpoint-rooted phylogeny of *L. borgpetersenii*, showing PathPhynder placement of the low-coverage ancient detection Kaz4. (C) *E. rhusiopathiae* phylogeny, rooted with an *E. tonsillarum* outgroup, showing PathPhynder placement of low-coverage ancient detections alongside the high-coverage ancient genome Galicia3. Alongside the tree, a tip-aligned heatmap shows per-virulence-gene read coverage (%), grouped into functional categories, for 26 genomes with available coverage data (modern strains and ancient genomes with mean coverage > 1); remaining tips are left blank (not analysed).

For *L. borgpetersenii*, we assembled a maximum likelihood phylogeny of 28 genomes, comprising 27 modern *L. borgpetersenii* references spanning isolates from cattle, humans, and small mammals, onto which we placed the low-coverage ancient sample Kaz4 (a medieval-era domestic cow from Georgia) using PathPhynder (Figure 3B). Modern genomes formed host-associated structure within the phylogeny, including a cattle-associated clade (largely serovar Hardjo-bovis–related isolates), a human clinical clade, and a rodent/small-mammal clade. Kaz4 fell within the cattle-associated *L. borgpetersenii* clade (Figure 3B), supporting the PIGSTI taxonomic assignment and consistent with recovery from an ancient *Bos* sample. The low number of reads recruited to the genome (n = 91 reads; ∼0.001× mean coverage) also precluded further downstream analysis of this strain.

*E. rhusiopathiae* was represented by a substantial number of ancient hits (n = 33) spanning a wide range of coverage (0.0007–67.71×). A high-coverage backbone phylogeny was built from 22 modern genomes, an *E. tonsillarum* outgroup, and the ancient genome Galicia3 (67.71×, recovered from a Neolithic aurochs sample from O Courel, Spain); this backbone separated the two marine-associated modern strains (dolphin, sea otter) from the remaining terrestrial diversity, within which Galicia3 branched basally. All 32 remaining low-coverage ancient hits placed successfully with PathPhynder, with most placements falling at basal nodes reflecting limited resolution at low coverage rather than distinct lineages (Figure 3C). Galicia1 placed directly with Galicia3, which have the same geographic and archaeological origin, rather than with any modern reference, indicating that shared geographic and temporal context can strongly shape phylogenetic placement of ancient *E. rhusiopathiae* strains.

Comparing virulence gene content among the four ancient genomes with ≥1× coverage (Galicia3, Dublin2, YG13, YG01) against 22 modern strains showed that variation was concentrated in a small number of surface-associated loci (Figure 3C), most consistently *ERH_0278*, *ERH_1258*, *ERH_1472*, and *ERH_1436*, which were reduced or absent in Dublin2, YG13, and YG01 and were also independently lost in the two most host-divergent modern strains (muskox, sea otter). Galicia3 showed recurrent loss of *ERH_0278* and *ERH_1472*. Beyond the surface proteome, Dublin2 also lacked the lysophospholipase C gene *ERH_1433* (mirroring its absence in dolphin, sea otter, and one domestic goose strain), while the biofilm-associated gene *ERH_1467* showed uniformly reduced coverage across both ancient and modern genomes in our dataset.

## Discussion

PIGSTI is a modular, reproducible Snakemake pipeline that combines host identification, metagenomic screening and pathogen authentication for ancient animal material. Existing workflows such as EAGER and aMeta provide established approaches for pathogen screening but do not incorporate host identification, which is particularly important when archaeological specimens have uncertain taxonomic assignments. PIGSTI integrates host identification with pathogen screening and applies between 10–13 metrics to evaluate candidate hits. In the empirical dataset, the initial Guellil et al. (*72*) screening approach identified 1,023 candidate pathogen hits, of which 109 passed the full authentication procedure. Phylogenetic placement of 36 genomes from three selected pathogens provided further support for these taxonomic assignments.

PIGSTI identified ancient pathogen-associated reads in 89 of 952 animal samples (9.3%), comparable to rates reported in other studies - for example, 15.9% as reported by Runge and colleagues (*54*), who particularly focused on pathological materials. In comparison, large-scale screening of human remains (*80*) found 52% with evidence of disease-associated microbial DNA. The lower rates of pathogen detection in faunal palaeogenomic datasets at least partially can be explained by the tendency for animal remains to be from food waste, and thus less likely to have died due to disease (*56*).

We also find tentative evidence that detections were distributed non-randomly across host species and sample materials. Enrichment of dental tissues (×2.7) is expected: teeth are well-established substrates for blood-borne and systemic pathogens (*119*). The host distribution is more surprising. *Ovis* accounted for 42% of positive specimens despite representing only 21% of the screened cohort (×2.0), an enrichment that persisted after excluding two datasets reporting only pathogen-positive data (*49*, *50*, *74*). An excess of pathogen detections from *Ovis* specimens may also appear in other studies, with Runge and colleagues finding twice as many detections in *Ovis* specimens (64% of all pathogen-positive specimens) as they represent in the dataset (32%). It is possible that these two datasets genuinely reflect a difference of infection rates between different species in the past, and that pathogen transmission was particularly common in sheep due to, for example, poor health conditions, denser herds, reduced genetic diversity due to intensive breeding, or greater exchange of sheep. However, we cannot exclude technical or sampling biases underlying this observation, such as higher DNA survival in sheep-rich assemblages, a larger number of *Ovis* among teeth samples, or greater sampling effort spent on well-preserved or pathological sheep specimens - all potentially biasing towards pathogen detection in such datasets.

The biological significance of the pathogen detections reported extends beyond taxonomic identification. Zoonoses account for a large share of the global infectious disease burden, and many pathogens that now infect humans circulated in animal populations before crossing the species barrier. Animal remains can therefore reveal pathogens circulating in livestock populations that are poorly represented in human palaeogenomic datasets, providing evidence for past pathogen diversity, animal infection and potential zoonotic exchange that cannot be reconstructed from human remains alone (*120*).

The two *R. felis* genomes illustrate this particularly well. STK240 and MC72713 were placed on the branch leading to *R. felis*, together with the previously reported ancient BBayA strain, supporting their assignment despite the low number of reads recruited. One dates to ∼9,000 BP, extending the known history of the pathogen into the Iranian Neolithic. Sikora et al. (*80*) did not detect *R. felis* in their large-scale screening of ancient human remains, despite recovering other vector-borne pathogens such as *Borrelia recurrentis*. Its detection here in Iran and Tunisia, together with the previous South African record, may therefore indicate a geographically restricted distribution that is poorly represented in human palaeogenomic datasets. The presence of *R. felis* in ancient livestock also raises the possibility of past human exposure through animal-associated vectors or contact.

*L. borgpetersenii* from a Georgian cattle specimen provides a different example. Its placement within cattle-associated *L. borgpetersenii* diversity, distinct from human clinical and rodent or small-mammal lineages, is consistent with its association with bovine leptospirosis. Its detection in the Caucasus at ∼1,200 BP falls within the period spanned by previously reported ancient human infections from five Eurasian samples dating between 5,000 and 900 BP (*80*). This does not establish a direction of transmission, but shows how combining animal and human pathogen genomes can reveal its presence in potential reservoir populations that would remain invisible when sampling humans alone. A final such example comes from the zoonosis *E. rhusiopathiae*. We expand on the number of ancient animal-associated *E. rhusiopathiae* detections; following the same filtering criteria as de-Dios and colleagues (*55*), who reported 7 such detections at 0.01× coverage and 1% genome evenness, we report 20 detections spanning ten millennia and four host genera, and 33 using our filtering criteria (including three *Bos*-associated samples reported in (*55*).

The simulations exposed a detection limit. PIGSTI failed to identify *M. leprae* and *B. anthracis*, assigning *M. leprae* reads to *M. avium* at low confidence. This most likely reflects limited genomic discrimination from closely related taxa in the database: *B. anthracis*, for example, shares >99% average nucleotide identity with members of the soil-associated *B. cereus* group (*121*). Applying thresholds at the species level may compound this problem by dispersing genuine signals across closely related taxa. Such pathogens therefore require complementary, genus-or clade-specific strategies, such as targeted alignment to diagnostic markers or reference panels designed to represent within-genus diversity, as applied to *Brucella* by L’Hôte and colleagues (*48*).

Reference bias represents a broader constraint. The NT-based database remains strongly weighted towards human-associated taxa (*122*), reducing the apparent representation of livestock-associated genomes and potentially missing genomic regions that are poorly represented among reference strains. Low coverage then further limits phylogenetic resolution: although 36 genomes could be placed, several contained too few informative sites to resolve finer relationships, particularly among *E. rhusiopathiae*. Expanding reference databases with livestock-associated and ancient genomes should improve taxonomic and phylogenetic resolution, while reference-independent assembly approaches could recover additional genomic information from well-covered samples without relying on the representation of individual reference strains (*123*).

## Conclusions

Ancient animal remains are an underused record of past infection, and the volume of published faunal genomic data is extending. PIGSTI makes that material tractable by doing in a single toolkit what has previously required separate workflows: it identifies the likely host species, maps and characterises the host genome, screens the remaining reads metagenomically, and authenticates candidate pathogens, so that host population genetics and pathogen detection are recovered together. Applied to 952 datasets it returned 109 authenticated detections across 89 samples, including the earliest *R. felis* yet reported, the first *L. borgpetersenii* from an ancient animal, and 20 newly detected *E. rhusiopathiae* genomes spanning 10,000 years and a range of host genera. Both livestock-specific pathogens and zoonoses were recovered, giving a window onto diseases that affected animal populations in the past and onto the animals that may have acted as reservoirs for human infection.

### Description of archaeological Sites for newly reported materials

#### Georgia

<u>Dariali</u> or <u>Tamara’s Fort</u> (42.738050, 44.625633) is located in Georgia, in the Greater Caucasus mountainous range, at an altitude of 1,350 m asl. The site was investigated between 2013 and 2016 within the ERC project “Persia and its Neighbours”, directed by E. Sauer, K. Pitskhelauri and others. The site bears evidence of long-term occupation spanning more than a millennium. The earliest evidence of human occupation at Dariali dates back to between the 6th and 4th c. BCE. The fort was constructed in the late 4th c. CE (or early 5th c. CE at the latest). The fort was then occupied intensively until the late 10th/early 11th c. CE and reoccupied between the late 13th and early 15th c. CE. Except for the 20th c., when some military structures were positioned within the fort, there is no traceable post-medieval occupation (*124*). A very large number of faunal remains were found and were studied by Mashkour and colleagues (*125*).

#### Iran

<u>Ganj Dareh</u> is an aceramic Neolithic mound located in the Kermanshah province of western Iran close between the city of Harsin and the Bisitoun massif. It was first excavated by Philip Smith in the late 1960s and 1970s and re-excavated by a joint Iranian-Danish mission in 2016 and 2017. The site comprises five major occupation levels dated to between c. 8200 - 7600 cal BCE. The most prominent is Level D, which preserved a dense cluster of rectangular structures. The faunal material from the most recent excavations are currently curated at the Department of Cross-Cultural and Regional Studies at the University of Copenhagen.

<u>Kuran Buzan</u> is located at Kuran Buzan Mountain, Hulailan Valley, Iran. The site was excavated by the Danish Archaeological Expedition in 1964. Faunal remains are from a habitation site on Kuran Buzan Mountain. The assemblage is of probable Bronze Age origin; the collection is at the Natural History Museum of Denmark (ZMK 115b/1965).

<u>Tang-i-Hamamlan</u>, is located at Kuran Buzan Mountain, Hulailan Valley, Iran. Excavated by the Danish Archaeological Expedition in 1964. Faunal remains are from a gorge or ravine on the mountain. Probable Bronze Age origin; the collection is at the Natural History Museum of Denmark (ZMK 115c/1965).

<u>Tang-e-ktcheck</u> is located at Kuran Buzan Mountain, Hulailan Valley, Iran. Excavated by the Danish Archaeological Expedition in 1964. Faunal remains from a cave habitation on Kuran Buzan Mountain. Probable Bronze Age origin; collection is at the Natural History Museum of Denmark (ZMK 115d/1965).

<u>Tepe Jarali</u> is located at Sar-i-Tarhan Valley, Iran. Excavated by the Danish Archaeological Expedition in 1964. Faunal remains from a house ruin on a small hill in the valley. These probably derived from the late Iron Age to Bronze Age; collection is at the Natural History Museum of Denmark (ZMK 115a/1965).

#### Uzbekistan

<u>Bukhara</u> is the principal urban centre of the oasis of the same name (in Uzbekistan) and has served as its political and economic capital since the Early Islamic period. The city consists of three main components: the Ark (citadel), the shahrestan (lower city), and the surrounding suburbs. Archaeological evidence indicates that the earliest occupation of the site dates to the 3rd century BCE. Initially organised around the citadel and an adjacent shahrestan located to the east, the settlement expanded considerably during the first centuries CE. Urban growth continued throughout Late Antiquity and the Early Medieval period, reaching a first apogee during the Samanid period (9th–10th centuries CE), when Bukhara became one of the major political and cultural centres of Central Asia. The city continued to expand during the Timurid period (14th–15th centuries CE) until the main urban transformation in the 16th and 17th centuries. Archaeological investigations conducted by the Franco-Uzbek Archaeological Mission in the Bukhara Oasis (MAFOUB) included a stratigraphic trench within the Ark (Trench A) and larger excavation areas in both the citadel (Trench C) and the shahrestan (Trenches B and D). The three specimens analysed in the present study (LL155, LL158, and LL162) all originate from Trench A within the Ark. Samples LL155 and LL158 were recovered from Islamic occupation layers dated to the 10th–12th centuries CE. Sample LL162 derives from a mixed archaeological context containing ceramic material dated between the 4th and 9th centuries CE. (*126*, *127*)

<u>Iskijkat</u> is situated in the central part of the Bukhara Oasis (Uzbekistan) on the route connecting Samarkand to Bukhara. The settlement follows the characteristic tripartite layout of many Central Asian urban centres, comprising a fortified citadel, an open shahrestan, and an extensive suburban area. Archaeological investigations by the Franco-Uzbek Archaeological Mission in the Bukhara Oasis (MAFOUB) included two deep stratigraphic trenches, one excavated within the citadel (Trench A) and the other in the shahrestan (Trench B). The earliest evidence of occupation dates to the 3rd–2nd centuries BCE. Subsequent occupation phases are documented during the 2nd–1st centuries BCE and again in the 2nd–3rd centuries CE. From Late Antiquity onwards, the settlement experienced a long period of continuity, with both the citadel and the surrounding urban areas remaining occupied between the 4th–5th and 7th centuries CE. During the 7th–8th centuries CE, occupation expanded beyond the core urban areas, suggesting renewed demographic growth. In the Early Islamic period (9th–10th centuries CE), settlement activity became increasingly concentrated within the citadel and shahrestan, while the suburban zones appear to have been primarily devoted to commercial and craft activities. Occupation contracted substantially after this period, and evidence for the 12th–16th centuries CE is currently restricted to the citadel, indicating a marked reduction in the extent of the settlement, particularly after the 13th century. The two specimens analysed in the present study originate from Iskijkat. Sample LL155 derives from a context dated between the 1st century BCE and the 2nd century CE, while sample LL156 was recovered from deposits attributed to a broader phase spanning the 2nd/3rd to the 6th centuries CE (*128–130*).

<u>Kakishtuvan</u> is located on the north-western edge of the Bukhara Oasis (Uzbekistan), along the corridor connecting Bukhara to Khorezm. Like several major urban centres of the oasis, the settlement displays a tripartite organisation composed of a fortified citadel, a shahrestan (lower city), and an extensive suburban area. Archaeological investigations have shown that the earliest occupation was established within the citadel during the 1st century CE, as documented by the stratigraphic sequence excavated in Trench A by the Franco-Uzbek Archaeological Mission in the Bukhara Oasis (MAFOUB). Available archaeological evidence indicates that the shahrestan developed later, probably during the 3rd–4th centuries CE. Settlement activity intensified during the 5th century CE, as reflected by an increase in both occupation and construction phases. The stratigraphic sequence suggests a gradual decline of the citadel during the 6th–7th centuries CE, followed by progressive abandonment in the decades after the Islamic conquest. Nevertheless, the site remained occupied during the 9th–12th centuries CE, with more limited evidence of activity continuing into the Timurid period. The specimen analysed in the present study (LL161) derives from a context dated between the 4th and 6th centuries CE (*126*, *127*, *130*).

<u>Kafir-kala</u> is located approximately 12 km southeast of Afrasiab (ancient Samarkand, Uzbekistan), along the Dargom canal. Covering around 20 ha, the site occupies a naturally defended position surrounded by watercourses and rises prominently above the surrounding plain. The settlement consists of a fortified citadel, a shahristan protected by moats and defensive earthworks, and an associated suburban area. Archaeological investigations conducted by the Uzbek-Japanese expedition initially focused on the citadel (Trench 1) before expanding to several areas of the shahristan. Excavations have revealed substantial architectural remains, including a large hall in the Sharestan Area (Trench 5). The earliest securely dated occupation of the site dates to the 4th century CE. Kafir-kala experienced significant development between the 5th and early 8th centuries CE before undergoing a major destruction episode during the early Islamic conquest. Archaeological evidence indicates subsequent reoccupation which ended abruptly with another destructive fire around the mid-8th century, while occupation continued intermittently through the Medieval period. Three specimens analysed in the present study originate from Kafir-kala. Samples LL159 and LL160 were recovered from 8th–9th century CE occupation levels within the hall located in the shahristan. Sample LL163 derives from Room 12 of the citadel and is likewise dated to the 8th–9th centuries CE (*131–133*).

## Data and software availability

### Underlying data

Raw fastq files of the newly produced samples are available at ENA accession: PRJEB124647. This project contains raw shotgun sequencing data for the 153 libraries newly generated in this study. Accession numbers for individual libraries are given in Table S8 of the extended data.

### Extended data

Zenodo: Supplementary data and modified pathPhynder code for: PIGSTI, a modular, reproducible pipeline for detecting species identity, pathogens, and microbes from animal palaeogenomic data. Available at: : https://doi.org/10.5281/zenodo.22135224

This project contains the following extended data:

(1) Large_tables.xlsx, a workbook containing eight sheets: S1, the pathogen panel screened, with per-pathogen E-value and read-count thresholds; S2, sequencing data for the datasets newly generated in this study, giving for each library the sample of origin, library identifier, UDG treatment and number of raw read pairs; S3, detection scores and authentication metrics for the 60 simulated benchmarking datasets, across ten pathogens and six abundance tiers; S4, metadata for all 952 screened datasets, including sample identifiers, accessions, host taxon, material, age and provenance; S5, all 1,023 candidate hits passing the initial E-value threshold, with screening, mapping and authentication metrics; S6, the 109 hits passing the high-confidence filter set; S7, the 89 samples carrying at least one authenticated detection; and S8, metadata for the 153 datasets newly generated in this study, including archaeological labels and internal laboratory codes.
(2) Pathphynder_v1.2.3_modif: modified pathPhynder v1.2.3 implementing the --maximumToleranceProp option used for phylogenetic placement, with patch file and licence.

Extended data are available under the terms of the Creative Commons Attribution 4.0 International licence (CC-BY 4.0).

## Software availability

Source code available from: https://github.com/LouisLhote/PIGSTI. Archived source code at time of publication:https://doi.org/10.5281/zenodo.22161588. Licence: MIT. PIGSTI is implemented in Snakemake and requires Snakemake ≥ 7.32 and conda (*134*). Installation instructions and a test dataset are provided in the repository README. Modified pathPhynder v1.2.3 is archived in the extended data under the MIT licence, inherited from upstream pathPhynder (*92*).

## Acknowledgments

We thank Logan Kistler and Torben Rick for facilitating sampling at The Smithsonian Institution.

This work was supported by Taighde Éireann—Research Ireland grant 21/PATH-S/9515 (T) (K.G.D.); European Research Council grant 101220382-HERDPATH (K.G.D.); European Research Council grant 885729-AncestralWeave (D.B.)

